# Virtual Screening and Elucidation of Putative Binding Mode for Small Molecule Antagonist of BCL2 BH4 Domain

**DOI:** 10.1101/2020.07.29.226308

**Authors:** Ireoluwa Yinka Joel, Temidayo Olamide Adigun, Ahmeedah Ololade Ajibola, Olukayode Olusola Bankole, Ugochukwu Okechukwu Ozojiofor, Ifelolu Adeseye Remi-Esan, Lateef Adegboyega Sulaimon

## Abstract

Evading apoptosis is a hallmark of cancer cells, therefore therapeutic strategies have been developed to induce cell death. BCL2 family protein governs the intrinsic pathway of cell death. Targeting the BH4 domain to modulate the anti-apoptosis activities of BCL2 protein has been established however, BDA366 is the only BH4 binding molecule to be reported. Virtually screening ~ 1,000,000 compounds 11 putative BH4 binding small molecules with binding affinity ~ −84kcal/mol to - 64kcal/mol resulted. Using QM-polarized docking, Induced-fit docking, and QM-MM optimization, a putative binding mode for the top 3 compounds is proposed: compound 139068 interactions with GLU13, MET16, LYS17, ASP31, and GLU42; compound 138967 interactions with ASP10, ARG12, GLU13, HIS20, MET16, and GLU42; compound 38831 interactions with ASP10, ARG12, GLU13, LYS17, and GLU42. MD simulations (NMA) data showed the binding of the three compounds to be stable with low eigenvalues. Electronic properties derived via DFT calculations suggest chemical reaction of the compounds be via electrophilic reactions.

## Introduction

The uncontrolled proliferation of cancer cells is not only driven by oncogenes but also results from defective apoptotic machinery or a combination of both [1][2] as well as the development of resistance by cancer cells to chemotherapeutics-induced apoptosis. These culminate in poor prognosis with an 18% overall survival rate in cancer patients [1]. Acquired resistance to apoptotic cell death is largely due to an over-expression of anti-apoptotic genes, and down-regulation or mutation of pro-apoptotic genes [3]. Therefore, overcoming resistance to apoptosis by activating apoptosis pathways has been a major focus in the development of therapeutic strategies for cancer treatment.

Apoptosis is an evolutionarily preserved mechanism of controlled cell deletion in nature playing a critical role such as the deletion of redundant or damaged cells in diverse fundamental body processes of multicellular organisms [4]. Its machinery consists of two major inextricably linked activation pathways: the extrinsic pathway via interaction of transmembrane receptors and death ligands (such as Fas, TNF-α, and tumor necrosis factor-related apoptosis-inducing ligand (TRAIL)) [5][6] and the intrinsic pathway with its consequential leakage of multiple pro-apoptotic proteins including cytochrome c and Smac/DIABLO from the mitochondria to the cytosol due to mitochondrial membrane potential loss [7] as well as a third, though less well-known, initiation pathway known as the intrinsic endoplasmic reticulum pathway[6].

The BCL2 family of proteins play major roles in the intrinsic pathway of apoptosis; the family is subdivided into two: pro-apoptosis (BH123: BAK, BAX and BH3-only proteins: BID, BIM, BAD, BIK, PUMA, and NOXA) and anti-apoptosis (BCL2 protein, BCL-xL, MCL1, BCL-W, BCL-B, and BCL2a1) [8]

Briefly, the BCL2 family of protein regulates apoptosis in the following ways: firstly in the absence of an apoptosis stimulus, BCL2 protein (anti-apoptosis) binds with BH123 proteins: BAK and BAX (pro-apoptosis), thereby preventing activation i.e. formation of homodimers on the mitochondrion membrane causing the release of caspase and activating the intrinsic apoptosis pathway [9]; however, when an apoptosis stimulus is received BH3-only proteins binds with BCL2, preventing it from interacting with BH123 protein, this process, therefore, activate the pro-apoptotic protein BAK and BAX [5][9].

The complex binding and interactions within the BCL2 family occur via their BCL2 alpha-helical homology (BH) domains [20]. There exist four BH domains (BH 1-4) of which the BH3 domain is responsible for activation and inactivation interactions of pro-apoptosis and anti-apoptosis BCL2 family proteins. Since overexpression of BCL2 protein has implicated in several cancerous tumors [10], the BH3 domain has been targeted for developing small molecule inhibitors of BCL2 protein (anti-apoptosis) to induced cell death in cancerous cells; inhibitors such as ABT-737, ABT-263, ABT-199, Disarib, etc.[8] have been developed with varying degrees of success [5][6]. However, none has been approved for use; this is primarily due to the highly conserved nature of BH3 domains among the BCL2 family [10]; the search for small molecules inhibitors of BCL2 protein has thus continued.

Several studies have suggested developing small molecule inhibitors targeting the BH4 domain of BCL2 protein as a potential therapeutic strategy [9][11][12]. Han *et al* [13] recently used BDA366 as proof of concept to show the therapeutic potential in targeting the BH4 domain of BCL2 protein. The BH4 domain in BCL2 protein is important in the regulation of the anti-apoptotic activity of BCL2 protein; it is also required in the interaction (heterodimerization) of BCL2 protein with pro-apoptosis protein BAX [10]. Furthermore, mutant BCL2 protein without the BH4 domain has been shown to promote apoptosis instead of inhibiting apoptosis [10][12].

At the time of writing, BDA366 is the only small molecule targeting the BH4 domain that has been reported. We, therefore, sought to virtually screen for potential BH4 binding small molecules and investigate a putative binding mode for the identified compounds.

## Materials and Methods

### 2.1. Small molecules Library

The L sample of the SCUBIDOO database [14] was downloaded for this study. The L sample which consists of ~ 1,000,000 compounds is a sample representation of the whole database which consists of ~21,000,000 compounds.

### 2.2. Highthrouput virtual screening

#### 2.2.1. Protein Preparation

Schrodinger‟s Protein preparation wizard [15] was used to import the BCL2 protein from PDB (PDB ID:1GM), and prepossessed: Hydrogen atoms were added, disulfide bonds were created, missing loops and side chains were added using Prime [16] Termini was capped, water molecules beyond 5Å from het groups were deleted, finally, het sates at pH 7.0 +/−2.0 was generated using Epik [4]. Hydrogen bonds in the protein structure were optimized, water molecule with less than 3 Hydrogen bonds to non-water residues/molecule was deleted, and the whole protein structure was minimized converging heavy atoms to RSMD: 0.3Å.

#### 2.2.1. Ligand Preparation

The Small molecule library was prepared using Ligprep [15]: the molecule was ionized generating all possible states at pH 7.0 +/−2.0, the ligand was desalted and tautomers was generated. Possible stereoisomers were also generated (50 per ligand).

#### 2.2.2. Active site Grid generation

Using the receptor grid generation grid of Schrodinger suit, the docking grid file was generated. However, since there is still no co-crystallized ligand in the crystal structure, we manually inputted the amino acid residues for the active sites based on information from the literature (AA 6-31) [10,13].

#### 2.2.3. Molecular docking

Using Schrodinger High throughput virtual screening (HTVS) workflow, the small molecule library was docked into the BH4 domain of BCL2 protein. The workflow included filtering the library based on drug and lead-likeness criteria, docking using the 3 different glide docking protocol (HTVS, SP, XP), and finally post-possessing using Molecular Mechanics-Generalized Born Surface Area (MM-GBSA) protocol.

#### 2.2.4. Molecular Mechanics-Generalized Born Surface Area

Molecular Mechanics-Generalized Born Surface Area (MM-GBSA) evaluated the binding free energies (binding affinity) and implemented geometric minimization of the docked protein-ligand complex [18]. The binding free energy was calculated using VSGB 2.0 implicit solvation model and OPLS-2005 via Prime [19]. The binding free energy is calculated using Eq.1

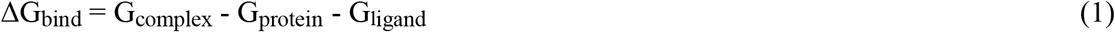

Where G_complex_, G_protein_, and G_ligand_ represent the free binding energy of the protein-ligand complex; protein; ligand respectively [18].

### 2.3. QM-Polarized Docking

QM-Polarized docking (qpld) module [20] of Schrodinger software was used to implement the qpld experiment. It involves three-step: firstly, docking using Glide SP [21] protocol; 50 poses per ligand generated at the of this stage. Secondly, Quantum mechanics (QM) charges for the generated ligand poses (50 per ligand) is calculated using a semi-empirical method (Charge type: Coulson). Thirdly, redocking of ligands with new QM charges using Glide SP protocol; 20 poses per ligand is generated.

### 2.4. Induced-fit Docking

Induced-fit docking experiment predicted active site conformational changes and ligand binding interactions to the new conformations. Using Schrodinger induce-fit extended sampling docking protocol [22], ligands were docked flexibly into the active site using Glide SP docking protocol [21] with ring conformation sampled at an energy window of 2.5kcal/mol; the side chain of the active site residues was trimmed with receptor and ligand van der waals scaling at 0.80. The maximum number of poses per ligand was set to 50. Thereafter the generated poses were refined using Prime [16]: the residues within 5.0Å of a ligand pose were refined and side chains optimized. The refined structures within 30.0kcal/mol of the best structure and the top 20 structure of overall were redocked using the Glide SP docking protocol.

### 2.5. Molecular dynamics

Molecular dynamics simulation using normal model analysis (NMA) was implemented using the iMOD server (http://imods.chaconlab.org) [23]. The simulation investigated the stability of the Induced-fit docked protein complex and the deformability of the complex. The following were calculated; Deformability, B-factor, Eigenvalues, Variance, Co-variance map, and Elastic network for the docked complex.

### 2.6. QM/MM Optimization

The resulting Induced-fit docking complex was optimized using QM/MM calculations [24] implemented via the Schrodinger Qsite module. The ligand and side-chain residues interacting with the ligand were treated as the QM region, while the protein was treated as MM region. DFT-B3LYP and basics set 631G** level was used for the QM calculations. The MM region was treated using OPLS2005 and energy was minimized using the Truncated Newton Algorithm.

### 2.7. Electronic Properties

The following Quantum mechanics (QM) electronic properties/descriptors (Highest occupied molecular orbital, lowest unoccupied molecular orbital, Electrostatic surface potential) were calculated using the Schrodinger Jaguar Single point energy module [41]. The module used hybrid Density flow theory (DFT) with Becke„s three-parameter exchange potential, Lee-Yang-Parr correlation functional (B3LYP), and basis set 631G** level [25][26]. From this aforementioned descriptors the following descriptors were calculated [27];

- HOMO-LUMO gap= E_LUMO_ – E_HOMO_,
- ionization energy (I) = −E_HOMO_,
- electron affinity (A) = −E_LUMO_,
- global hardness (η) = (−E_HOMO_ + E_LUMO_)/2,
- chemical potential (μ) = (E_HOMO_ + E_LUMO_)/2.
- global electrophilicity power of a ligand as ω = μ2/2η (proposed by Parr et al. [28]).

## Results

### 1.1. High throughput virtual screening

Molecular docking is the most used technique in computer-aided drug discovery. It predicts the energetically favorable binding conformation of ligands in the protein active site [29]; this enables virtual screening of large databases of ligands. This technique was used to screen ~1,000,000 ligands for interactions with the BH4 domain of BCL2 protein; after screening the binding affinity of the ligands was scored and geometrically minimized using MM-GBSA. BDA366 which has been established as a selective BH4 interacting ligand [30] was used as the standard for this experiment.

The data showed binding affinity of the ligands ranged from ~ −74kcal/mol to −24kcal/mol, with BDA366 having a binding affinity of −45.39kcal/mol. The top ranked 11 compounds binding interactions were analyzed (Figure 1).

**Figure 1:**
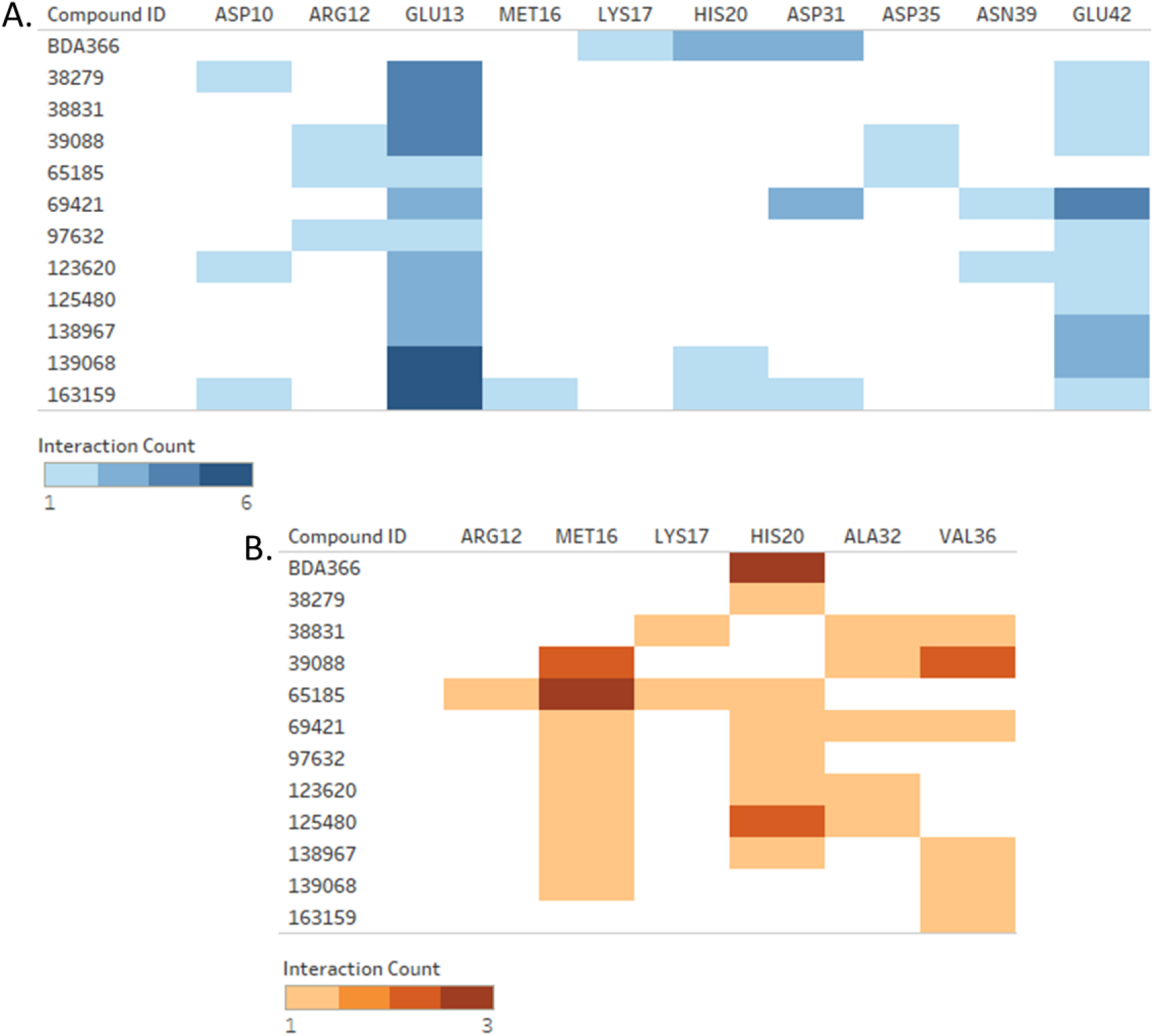
Molecular Docking binding interaction analysis. a) Hydrogen bond and electrostatic interaction count b) Hydrophobic Interaction Count

Visual inspection of BDA366 showed Hydrogen Bond (H-bond) interactions with LYS17, HIS20, and ASP31; electrostatic interactions with ASP31 (Attractive charge), and HIS20 (Pi-cation); hydrophobic interactions with HIS20 (pi-pi T-shaped) (Figure 2). The binding conformation of the first 3 compounds (125480: −74.67kcal/mol, 38279: −71.97kcal/mol, 139068:−67.22kcal/mol) was visualized and examined in detail (Figure 3). Compound 125489 formed H-bond interactions with GLU13 and GLU42; electrostatic interactions with GLU13 (Attractive charge); hydrophobic interactions with ALA32 (Alkyl), HIS20 (Pi-alkyl), Met16 (Alkyl) (Figure 3). Compound 38279 formed H-bond with ASP10 and GLU13; electrostatic interactions with GLU13 (Attractive charge) and GLU42 (Attractive charge); hydrophobic interactions with HIS20 (Alkyl) (Figure 3). Compound 139068 formed H-bond with GLU13 and GLU42; electrostatic interactions with GLU13 (Attractive charge), GLU42 (Attractive charge), and HIS20 (Attractive charge); hydrophobic interactions with VAL36 (Alkyl) and MET16 (Alkyl) (Figure 3). Generally, interaction with GLU13, HIS20, VAL36, and GLU42 was the most observed interactions among the 11 compounds (Figure 1).

**Figure 2:**
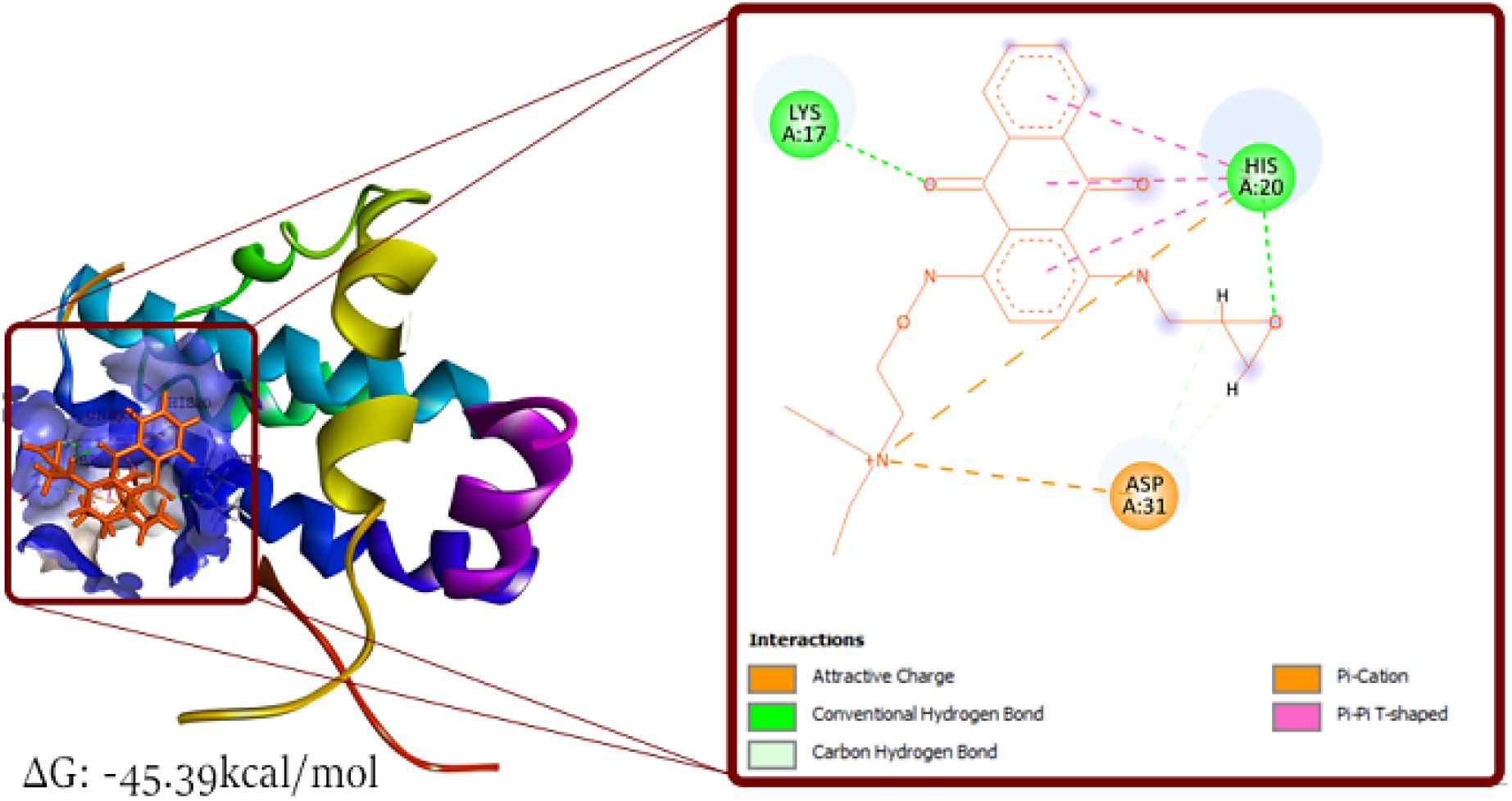
Molecular docking pose visualization: BDA366.

**Figure 3:**
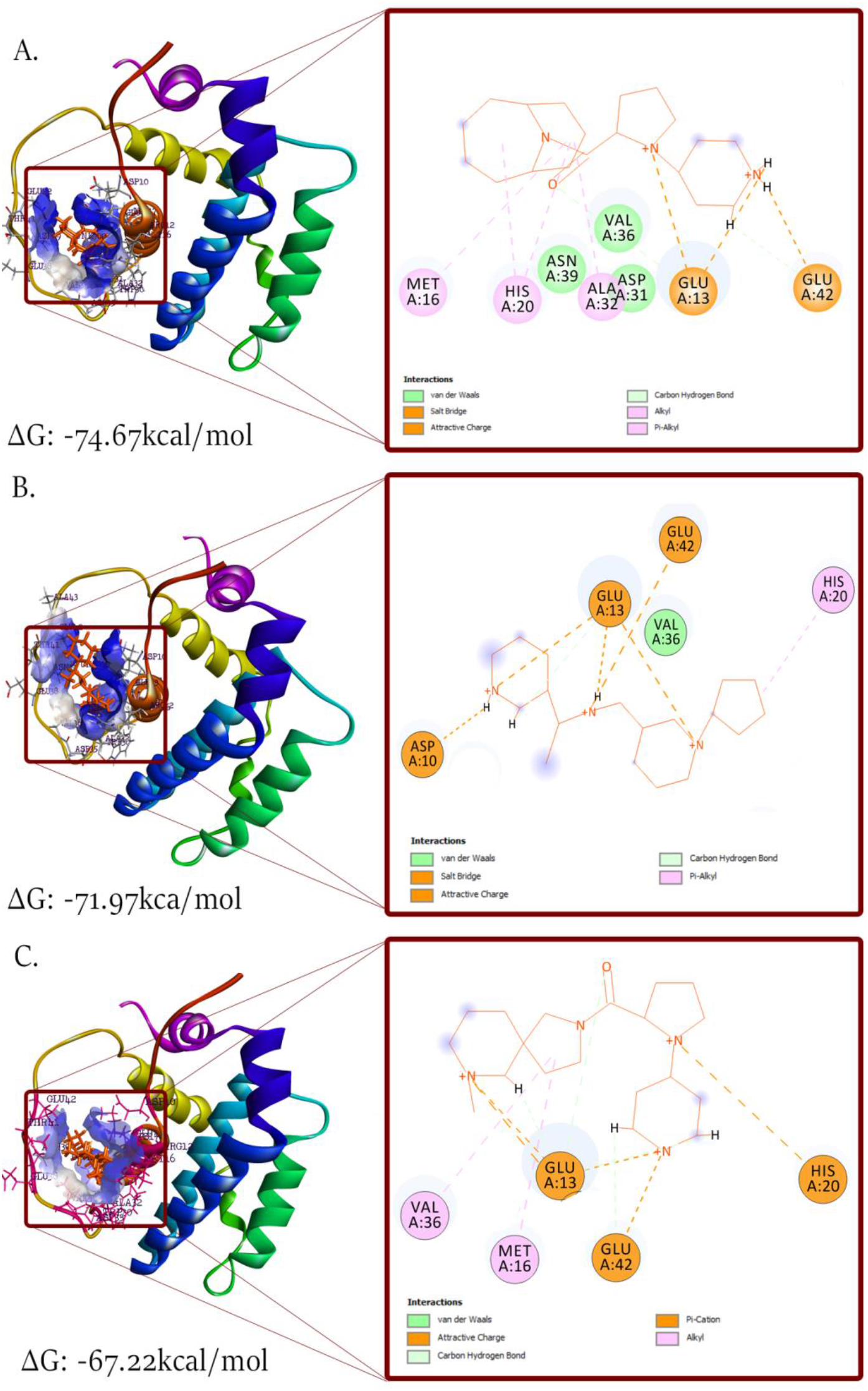
Molecular Docking pose visualization. a) 125480 b) 38279 c) 139068

Using molecular docking we were able to virtually screen ~1,000,000 compounds and identified 11 compounds interacting with the BH4 domain. Binding affinity ranged from −74 to −63kcal/mol, all having a higher binding affinity than BDA366 (−45.39kcal/mol). The next stage of the study was to elucidate a binding mode; the 11 compounds were hence subjected to more rigorous and accurate binding simulation protocols.

### 1.2. QM-Polarized Docking

Electrostatic charges on ligand atoms influence the accuracy of the simulation (binding affinity and binding interactions) [29]. Apart from initial atomic charges, charges induced on the ligands by the active site residues greatly improve docking simulation accuracy [20][28][31]. QM-polarized docking (qpld) uses this induced charge in its calculations (see methods).

Using BDA366 as the control for this docking run we docked the 11 compounds and calculated binding affinity using MMGBSA. The binding interaction analysis showed BDA366 to have acquired new H-bonds interactions with SER24, TYR21, ILE14, HIS94, and Hydrophobic interactions with LEU95 (pi-alkyl). When compared with the molecular docking binding interactions, a new hydrophobic interaction with LYS17 (pi-alkyl) is formed alongside an electrostatic interaction (pi-cation). The hydrophobic and electrostatic interactions with HIS20 were lost and only the H-bond retained. An increase in the binding affinity from −45.39kcal/mol to −49.89kcal/mol was observed (Figures 4 and 5).

**Figure 4:**
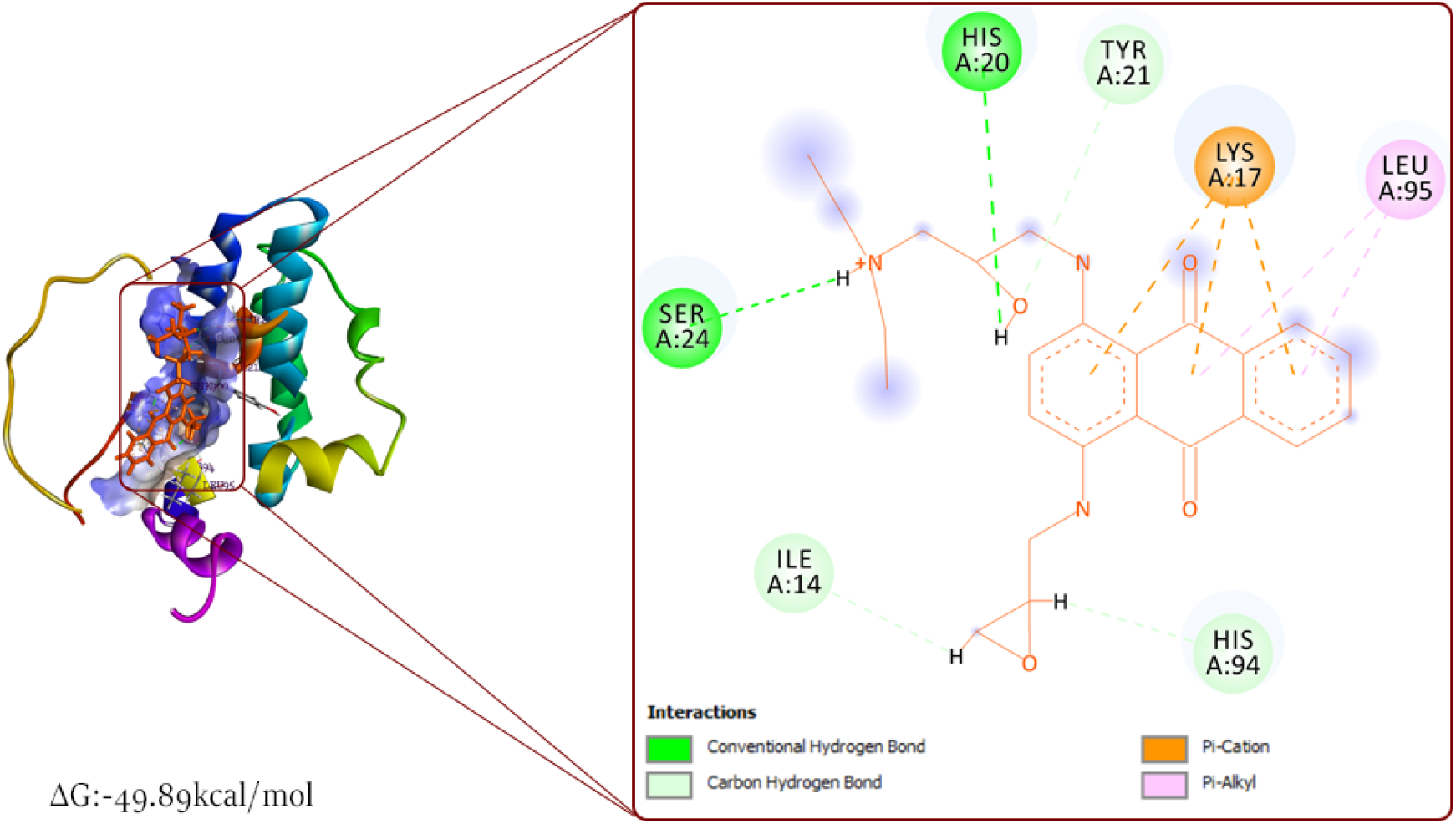
QM-Polarized Docking pose visualization: BDA366.

**Figure 5:**
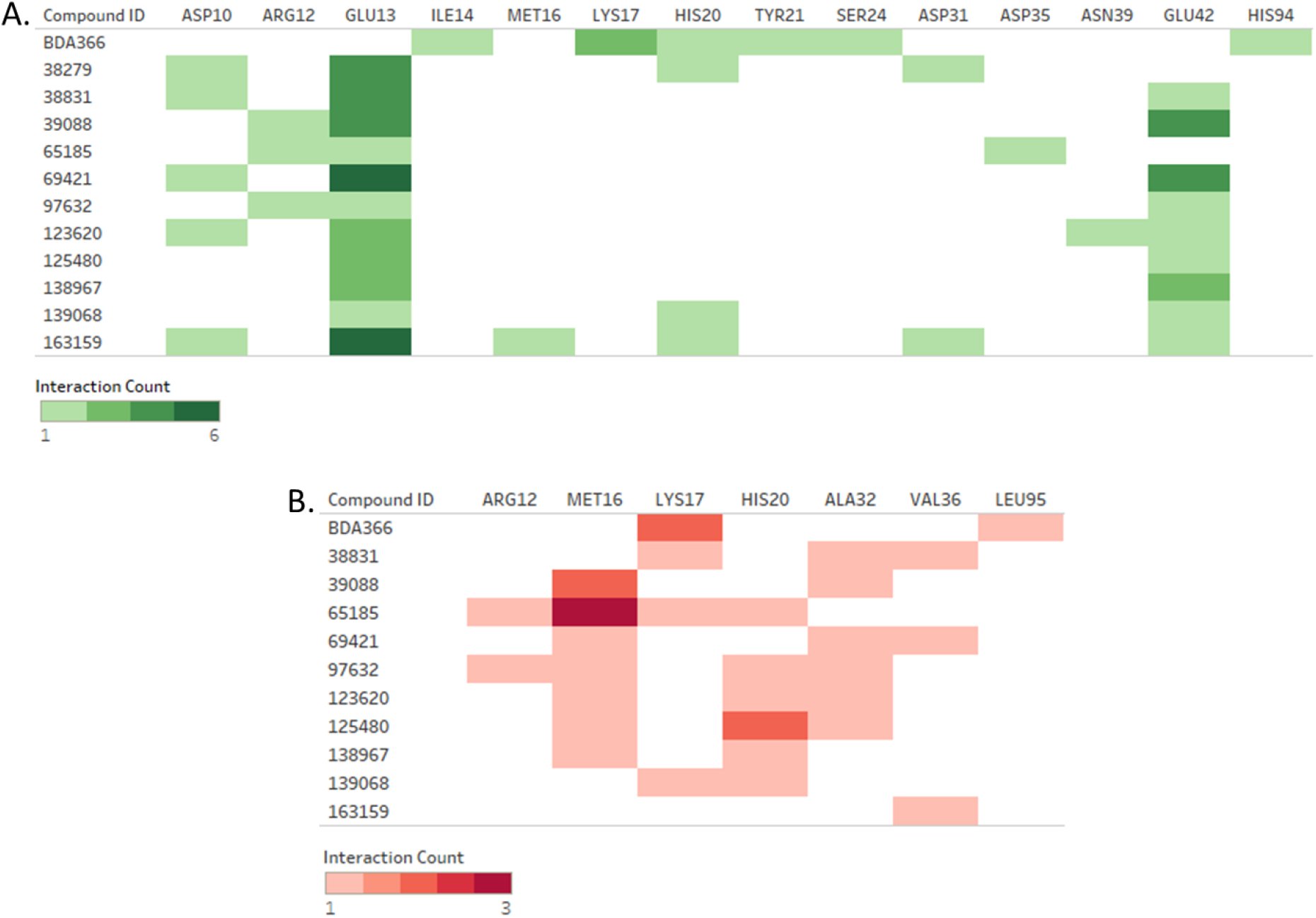
QM-Polarized Docking binding interaction analysis. a) Hydrogen bond and electrostatic interaction count b) Hydrophobic Interaction Count

The top three compounds were visualized (Figure 6). Compound 125480 had a binding affinity of 74.63kcal/mol. It formed H-bond with GLU42 and GLU13; an electrostatic interaction with GLU13 (attractive charge); hydrophobic interactions were formed with ALA32 (alkyl) and MET16 (alkyl) (Figure 5 and 6). When compared with the molecular docking binding pose, all interactions were retained and there was no significant change in binding affinity (−74.67 to −74.63kcal/mol). Compound 38831 had a binding affinity of −70.5kcal/mol, H-bond interactions were formed with GLU13 and GLU42; electrostatic interactions with ASP10 (attractive charge); Hydrophobic interactions with VAL36 (alkyl), LYS17 (alkyl), and ALA32 (alkyl) (figure 5 and 6). When compared with the molecular binding interactions it lost its H-bond formed with GLU42 and there was an increase in binding affinity from −63.81 to 70.95kcal/mol (Figures 1 and 6). Compound 38279 had a binding affinity of −67.76kcal/mol, it formed H-bond interaction with ASP10, GLU13, and ASP31; electrostatic interactions with GLU13 (attractive charge), ASP10 (attractive charge), and HIS20 (pi-cation) (figure 5 and 6). Comparing both docking pose (molecular docking and QM-polarized docking) we observed a lost electrostatic interaction with GLU13 (attractive charge) and also a hydrophobic interaction with HIS20 (figure 1 and 6). Also, there was a reduction in the binding affinity (−71.97 to −67.76kcal/mol).

**Figure 6:**
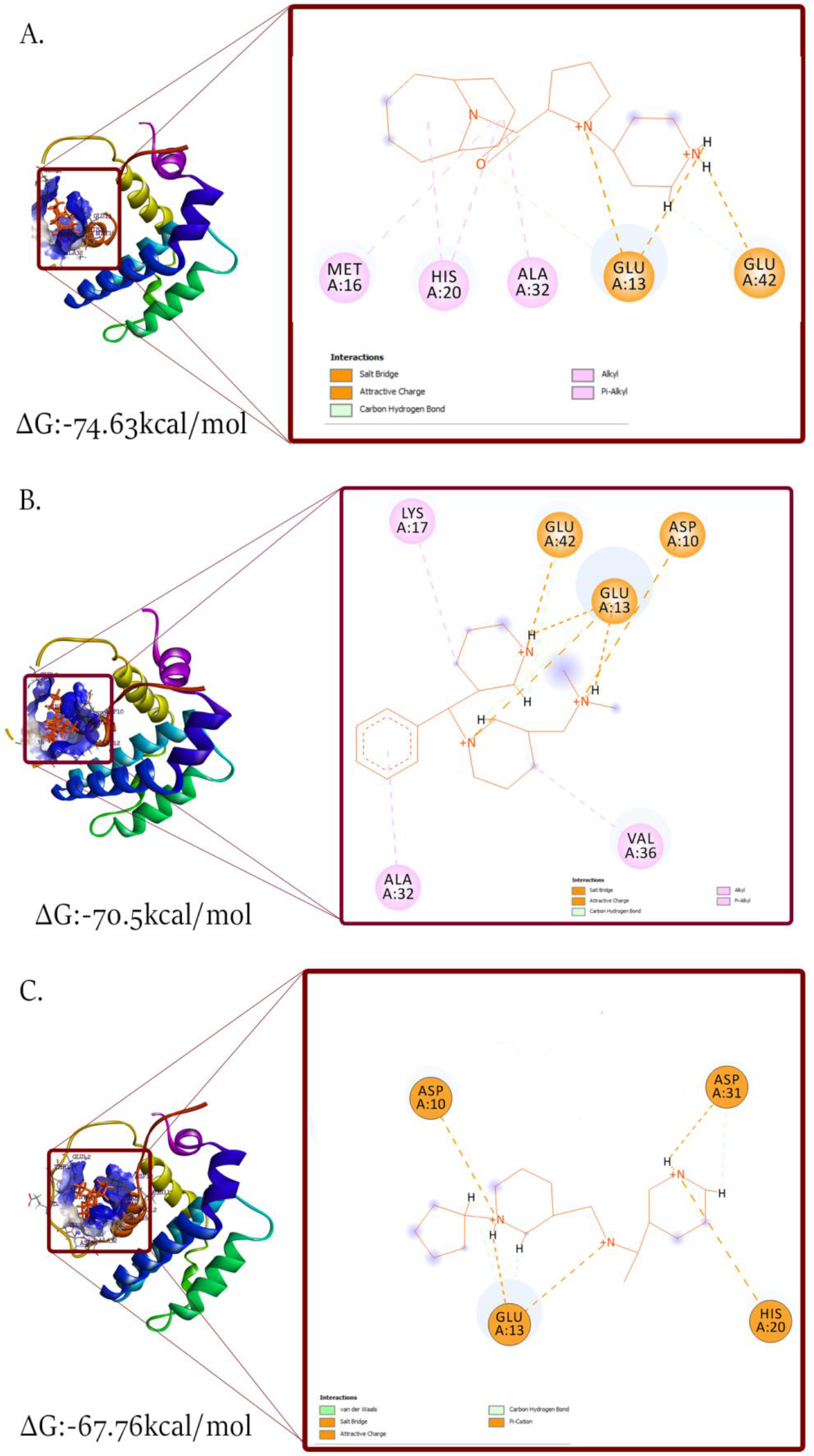
QM-Polarized Docking pose visualization. a) 125480 b) 38831 c) 38279

General analysis of the 11 compounds binding interactions showed the most common BH4 residue interactions were with GLY13, MET16, and HIS20 while non-BH4 residues: ALA32 and GLU42 (figure 5).

### 1.3. Induced-Fit Docking

In a biological system, both ligand and protein are flexible; the binding of a ligand induces series of conformation changes in the protein active site leading to a fit formation around the ligand. [32][22]. However, classical docking holds the protein active site rigid. Advances in docking techniques have allowed for the combination of rigid receptor-ligand docking protocol and protein structure prediction protocol to simulate and predict these conformational changes. This method has accurately predicted protein-ligand binding conformation of various protein targets [39][47][33]. Using Schrodinger‟s induced-fit docking module (see methods), the 11 compounds (BDA366 as control) were subjected to Induced fit docking and binding affinity calculated using MMGBSA.

Binding interaction analysis showed an increase in the number of interactions, with binding affinity increasing significantly generally (figure 7, 10). BDA366 formed H-bond with ASN11, ARG12, GLU13, MET16; electrostatic interaction with ASP10 (salt bridge), and GLU13 (pi-anion and attractive charge); hydrophobic interaction with ALA32 and VAL36 (figure 8). This data is consistent with the BH4 residues suggested by Han *et. al* [30]. It was suggested that BDA366 interacts with ASP10, ASN11, ARG12, and GLU13 in other to regulate BCL2. BDA366 had a binding affinity of −64.90kcal/mol.

**Figure 7:**
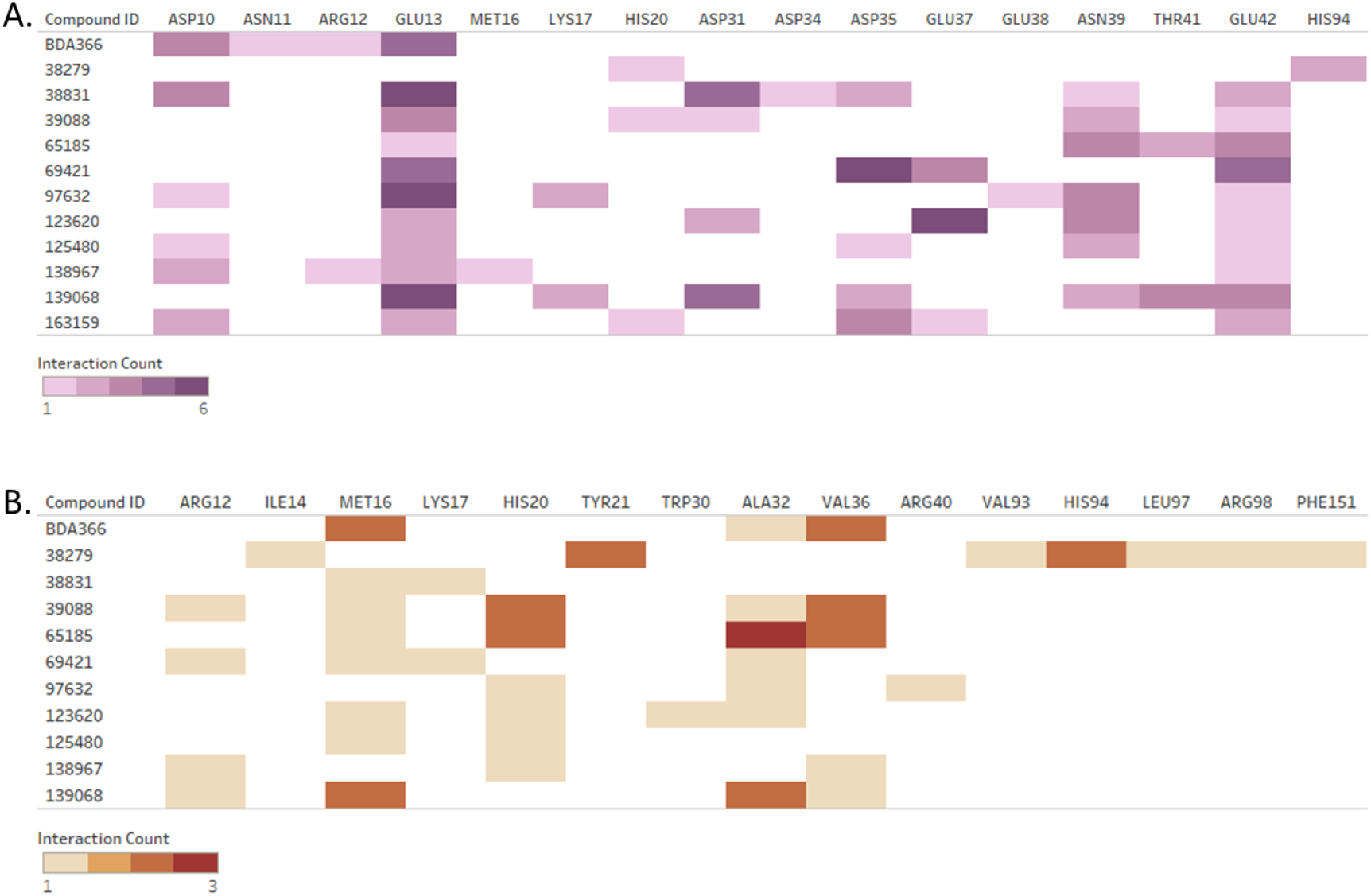
Induced-Fit Docking binding interaction analysis. a) Hydrogen bond and electrostatic interaction count b) Hydrophobic Interaction Count

**Figure 8:**
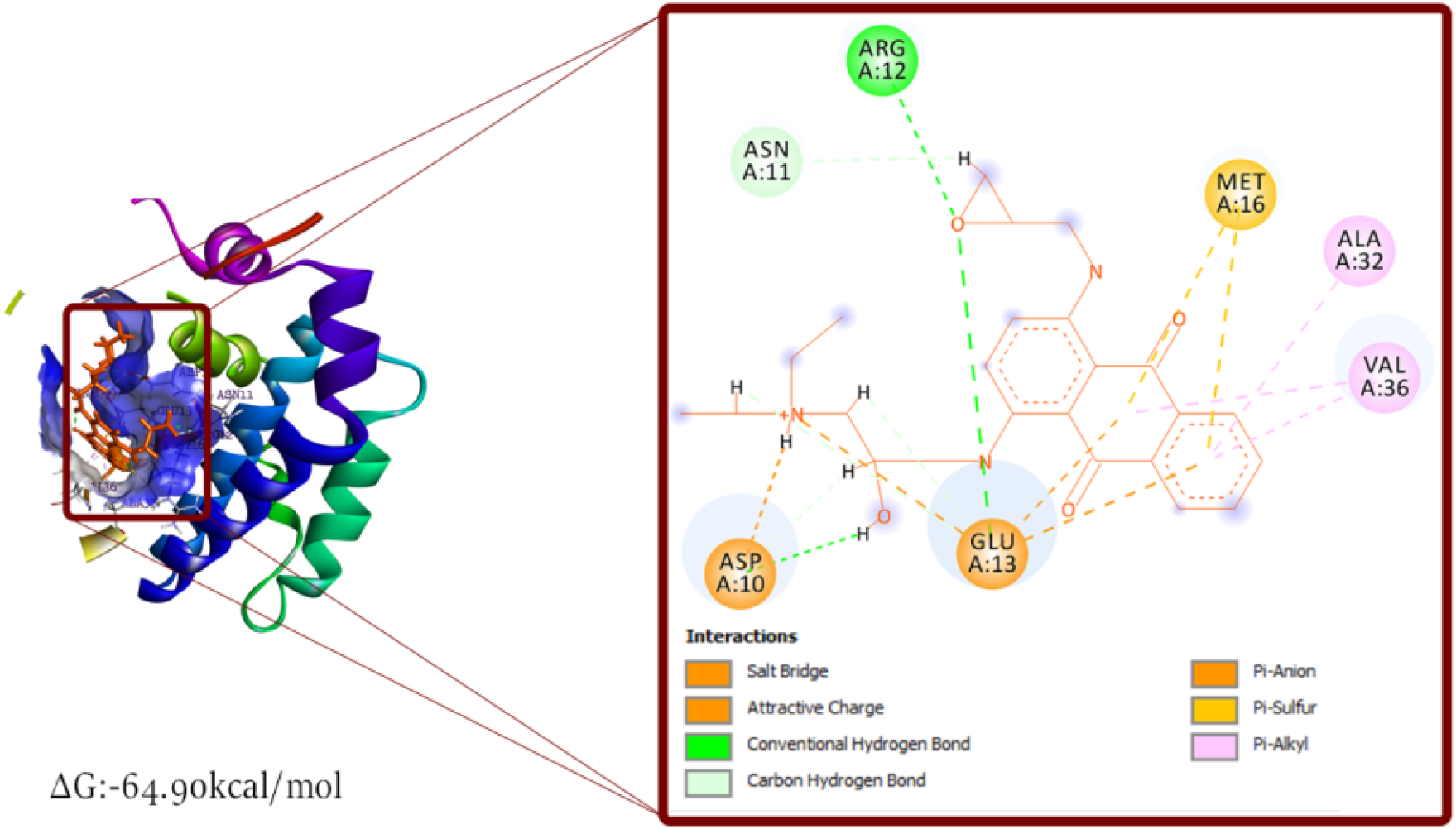
Induced-Fit Docking pose visualization: BDA366.

The 11 compounds had their binding affinity ranging from −100 to −85kcal/mol. The top 3 ranked compounds were 139068, 138967, and 38831 (−100.78kcal/mol, −95.27kcal/mol, and −93.20kcal/mol respectively.). Compound 139068 interacted with BH4 residues AA: 12, 13, 16, 17, and 31. Non-BH4 interactions included AA: 32, 32, 36, 41, and 42. Compound 138967 interacted with BH4 residues AA: 10, 12, 13, 16, 20, and non-BH4 residues AA: 36 and 42.

Compound 38831 interacted with BH4 domain residues AA: 10, 13, 16, 17, and 31 and non-BH4 domain residues AA: 35, 34, 39, and 42 (figures 7, 9).

**Figure 9:**
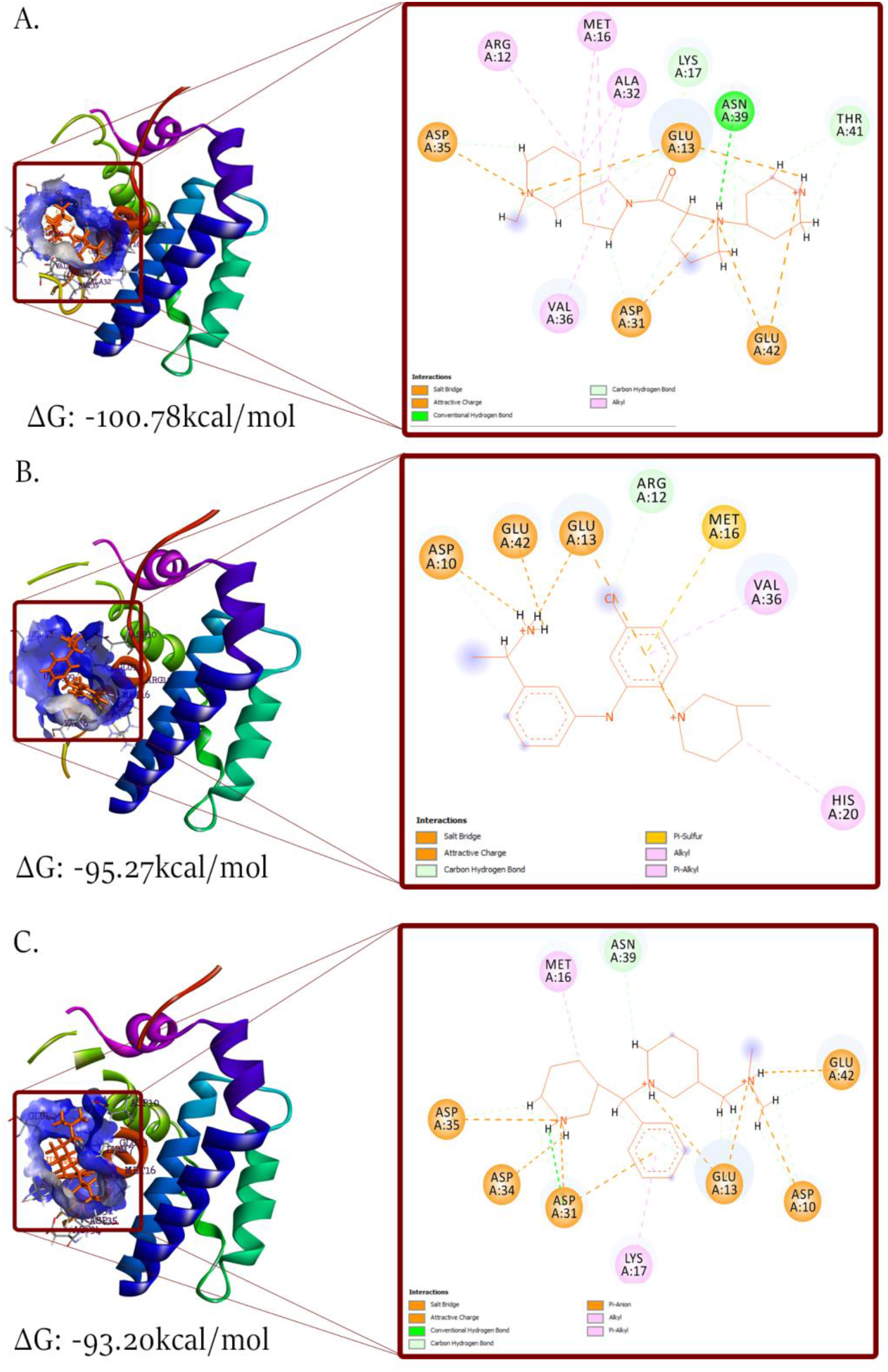
Induced Fit Docking pose visualization. a) 139068 b) 138967 c) 38831

In consolidating the binding affinities from the three different docking protocols a weighted average was calculated. The weights were calculated by adding a point (1) for every interaction with a BH4 residue (“reward”) and subtracting 0.5 for every interaction with non-BH4 residues (“punish”). We infer that ligand interactions with BH4 domain residues should be giving more priority and “rewarded” as the aim of this study is to identify BH4 binding molecules.

The weighted average binding affinity was calculated for the 11 compounds including BDA366 (Figure 10). Compound 139068, 138967, and 38831 were the top 3 compounds with the binding affinity of −84.46kcal/mol, −83.88kcal/mol, and −80.49kcal/mol respectively (figure 11). These compounds were selected as representative of the 11 for further analysis.

**Figure 10:**
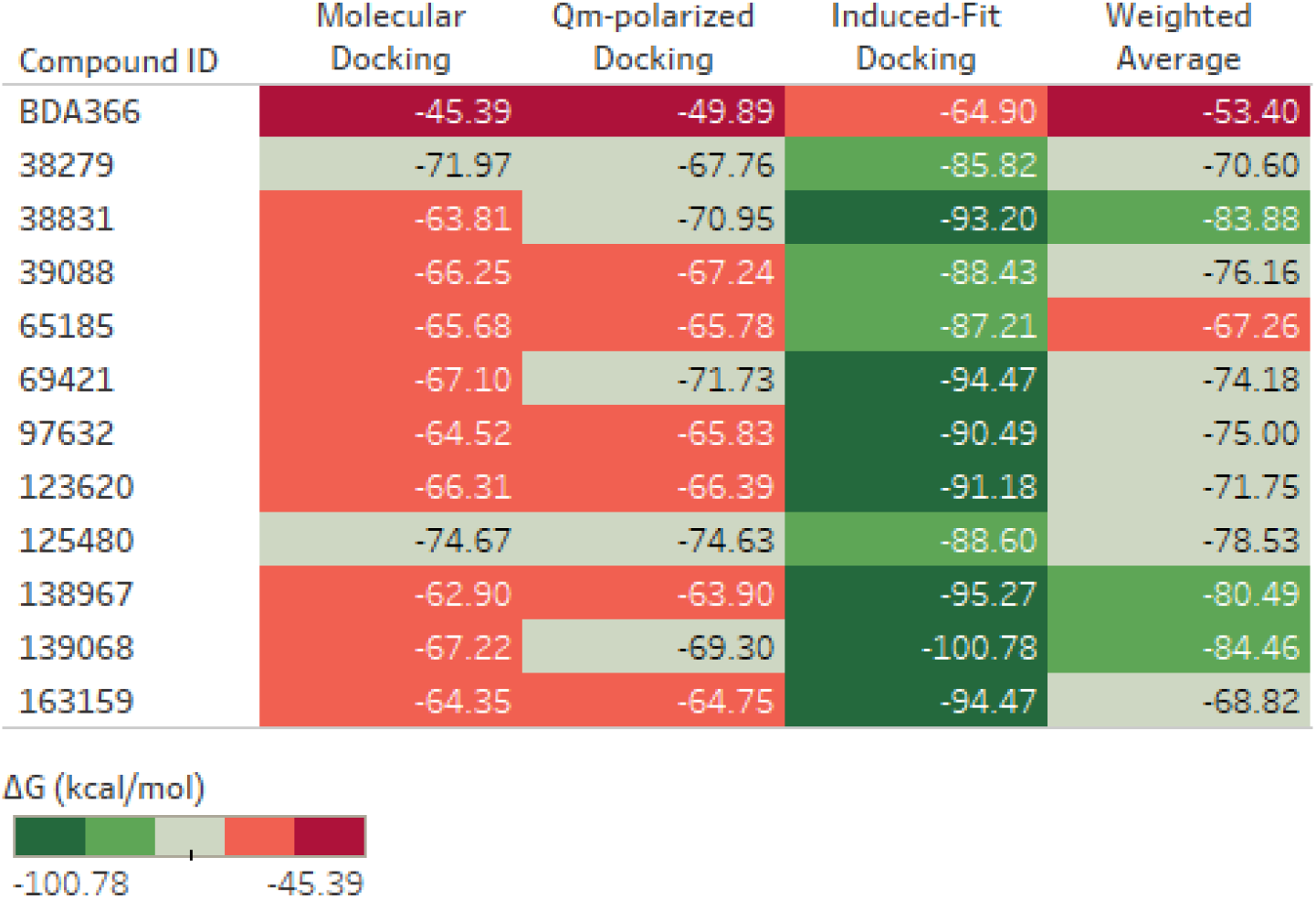
Ligand binding affinity for different Docking Protocol.

**Figure 11:**
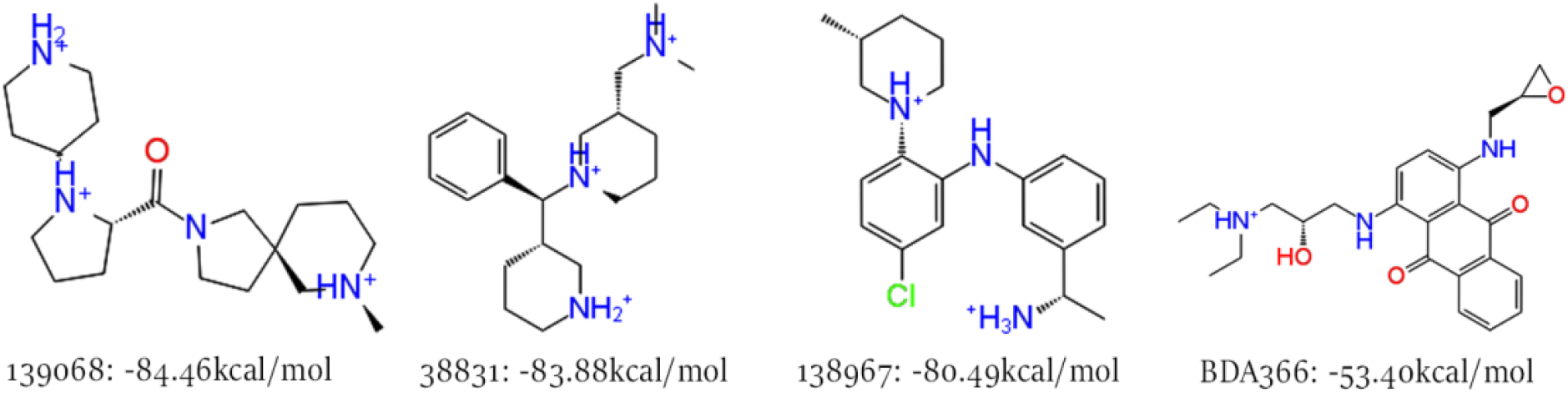
Top 3 representative compounds (based on weighted average) with BDA366(control)

### 1.4. Molecular Dynamics

Binding of BDA366 to the BH4 domain induces conformational changes leading to the exposure of BCL2 [30] BH3 domain. Using Normal mode analysis (NMA) we investigated the stability of the induced-fit docking complex and the deformability of the protein as a result of ligand binding. NMA investigates the flexibility of a molecule (protein) about an equilibrium position [34]. NMA assumes a single conformation for the docked complex and simulates an oscillatory vibration by connecting assumed springs between the residues forming an elastic network. The motion is broken into individual modes defined by eigenvectors and eigenvalues [35].

Eigenvalues represent the motion frequency of a mode, high eigenvalues represent low-frequency motions (important for enzyme catalysis) while low eigenvalues represent high-frequency motions (important for ligand binding and stability) [35]. Although not as accurate as classical Molecular dynamics (MD) NMA has been used in several studies to study protein dynamics (including ligand-induced conformational changes) [36][37][34][35] and has shown appreciable correlations with classical MD simulations and experimental data. Using iMOD server (see methods) NMA analysis was carried out.

The following were calculated: the deformability of the protein structure with peaks representing regions of high deformability (figure 12i), B-factor (figure 12ii), eigenvalues (figure 12iii), the variance associated to each normal mode which is inverse to the eigenvalues (figure 12iv), the covariance map indicating correlated (red), uncorrelated (white) and anti-correlated (blue) motion between pairs of residues (figure 12v) and finally, the elastic network representing the spring connections and their relative stiffness indicated by a color bar (figure 12vi). However, of uttermost importance is the eigenvalues calculated (figure 12iii; table 1).

**Table 1:** Molecular Dynamics Eigenvalue.

| Compound ID | Eigenvalues |
| --- | --- |
| 139068 | 2.62687e-04 |
| 138967 | 2.744942e-04 |
| 38831 | 2.591518e-04 |
| BDA366 | 2.682577e-04 |

**Figure 12:**
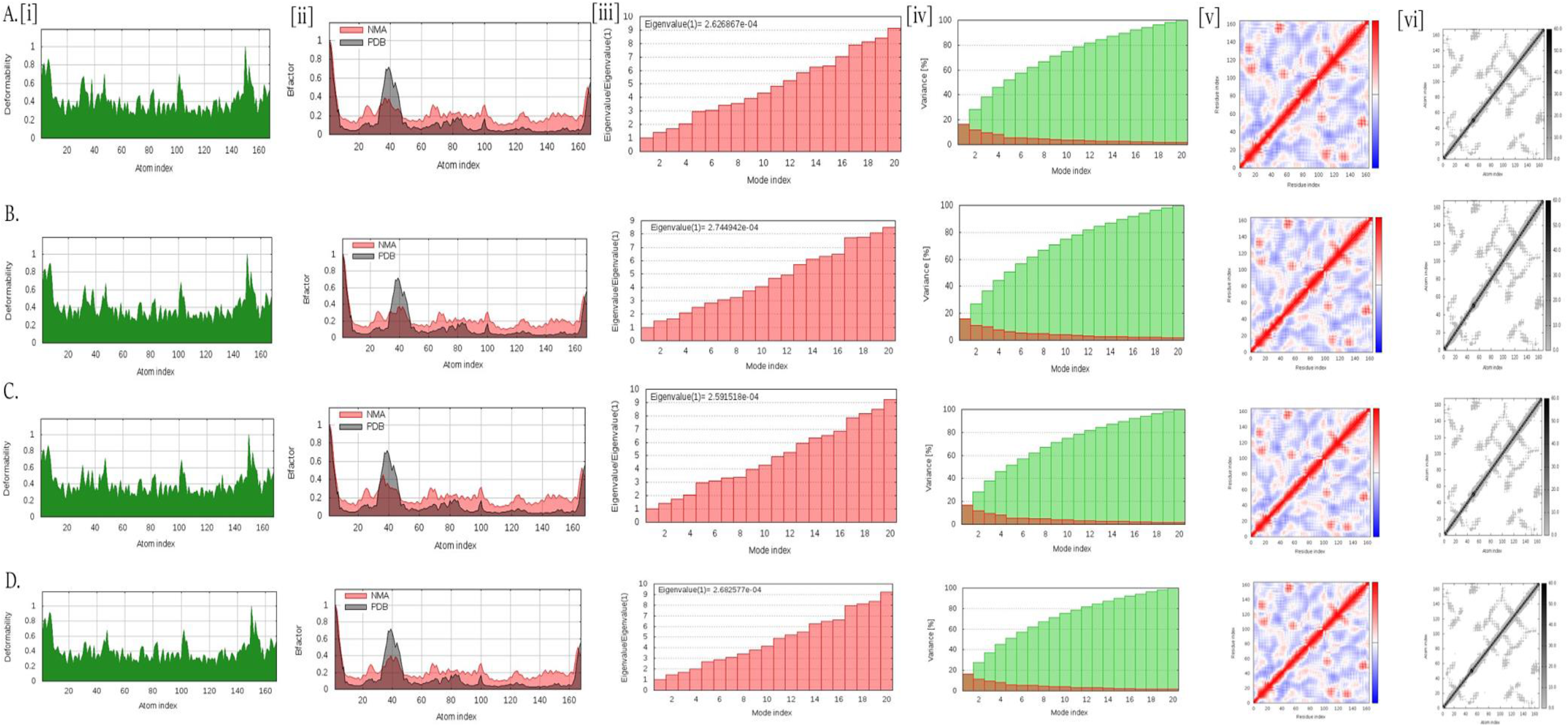
Normal mode analysis simulation of a) 139068 b) 138967 c) 38831 d) BDA366 induced-fit docking pose. i) Deformability ii) B-factor iii) Eigenvalues iv) Variance v) Co-variance correlation plot vi) Elastic network

As stated above eigenvalue is a function of individual mode motions, it represents motion stiffness of the protein and its value correlates with the energy required to deform a protein structure, therefore, lower values indicate the ease of deformation [23]. Compound 38831 had the lowest eigenvalue: 2.591518e-04 followed by 139068 (2.62687e-04) and 138967 (2.744942e-04) respectively (Table 1). BDA366 had an eigenvalue of 2.682577e-04. Generally, the 3 compounds and BDA366 showed low eigenvalue suggesting stable ligand binding and relative ease of deforming the protein complex.

### 1.5. QM-MM Optimization

The molecular dynamics investigated ligand binding stability and ease of deforming the protein complex (conformational changes), Quantum mechanics - Molecular mechanics (QM-MM) calculations were used to validate ligand binding interactions. The ligand and interacting side-chain residues were treated as the QM region while the protein structure as MM region. This calculation geometrically optimized the induce-fit docking complex and validated interactions observed. Using Qsite (see methods) the QM-MM optimization was implemented, after which the binding affinity of the optimized structure was calculated using MM-GBSA. The pre-optimized pose (IFD pose) and post-optimized pose (QM-MM pose) alongside their corresponding binding affinity were thereafter compared.

The optimized complex of compound 139068 maintained all interactions formed (when compared with pre-optimized pose) and a slight increase in calculated binding affinity (−102.15kcal/mol) (figure 13). Two new interactions were formed with Compound 138967: H-bond with GLU37 and electrostatic interactions with GLU38. H-bond with ARG12 was changed to hydrophobic interactions. This observed interaction changes resulted in the increase in binding affinity from −95.27 to −99.20kcal/mol (figure 13). Compound 38831 formed a new H-bond with THR41 and additional electrostatic interaction with ASP31. This new interaction however did not result in an increase in binding affinity but a reduction: −93.20 to −80.69kcal/mol (figure 13).

**Figure 13:**
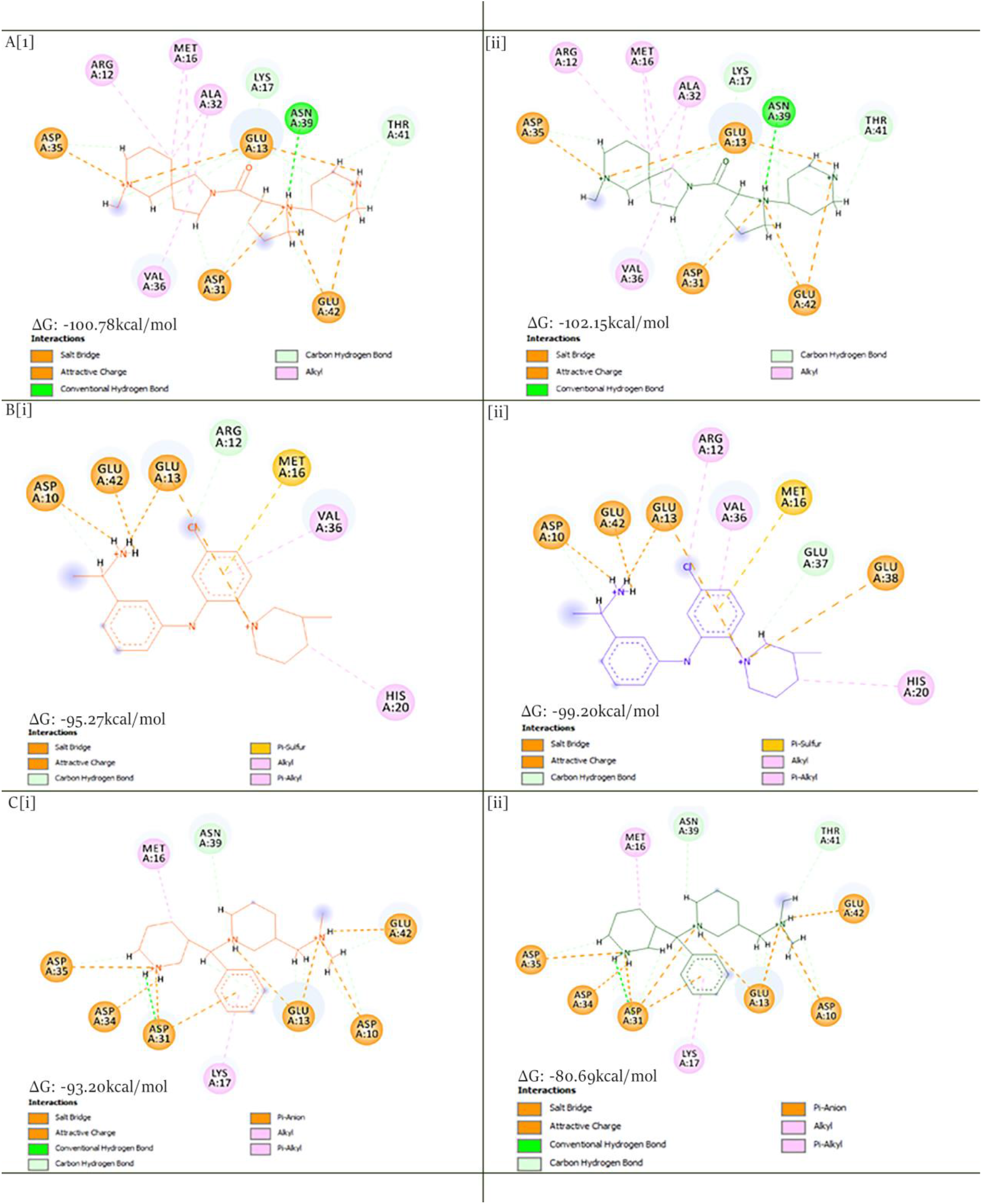
Comparison of pre (i) and post (ii) optimized induced-fit binding pose. a) 139068 b) 138967 c) 38831

### 1.6. Electronic Properties

Electronic property (descriptors) of a ligand provides insight into how a ligand might exercise its biological activity and provides insight on how to optimize for better biological activity. Using DFT calculations implemented via Single point energy module (see methods) electronic descriptors were calculated to investigate reactivity, mechanics of reaction (electrophilic or nucleophilic reaction), and stability of the 11 compounds. The descriptors calculated include HOMO, LUMO, and MESP; from these descriptors, the following were extrapolated: HOMO-LUMO gap, Ionization energy, Electron affinity, Chemical potential, Global hardness, and Global electrophilicity.

Highest occupied molecular orbital (HOMO), lowest unoccupied molecular orbital (LUMO) is one of the most important orbitals in a compound; calculating the HOMO-LUMO gap indicates ease of electron movement (from a region of LUMO to HOMO) and stability of the compounds; a small HOMO-LUMO gap indicates a more reactive compound but less stable compound [38][39]. Ionization energy is the energy required to remove an electron from a gaseous atom (it gives information about the energy of the orbital it originated from) while electron affinity is the energy change/released when an electron is added to a gaseous atom. Generally, electrons move from regions of high chemical potential to regions of low chemical potentials; therefore, a compound with low chemical potential indicates an electrophile. The Global hardness of a compound corresponds to the HOMO-LUMO gap of the compound [49]; hardness indicates the strength of electrophilicity. Global electrophilicity indicates the general reactivity of the compound. These descriptors are shown in Table 2.

**Table 2:** Quantum electronic descriptors.

| Compound Id | HOMO (eV) | LUMO (eV) | HOMO-LUMO GAP | Ionization energy (I) | Electron Affinity (A) | Global Hardness ($\eta$) | Chemical Potential ($\mu$) | Global Electrophilicity ($\omega$) |
| --- | --- | --- | --- | --- | --- | --- | --- | --- |
| **BDA366** | -0.26 | -0.17 | 0.09 | 0.26 | 0.17 | 0.09 | -0.22 | -2.39 |
| **139068** | -0.57 | -0.36 | 0.21 | 0.57 | 0.36 | 0.21 | -0.47 | -2.21 |
| **138967** | -0.43 | -0.25 | 0.18 | 0.43 | 0.25 | 0.18 | -0.34 | -1.89 |
| **163159** | -0.6 | -0.32 | 0.28 | 0.6 | 0.32 | 0.28 | -0.46 | -1.64 |
| **38831** | -0.54 | -0.33 | 0.21 | 0.54 | 0.33 | 0.21 | -0.44 | -2.07 |
| **123620** | -0.39 | -0.24 | 0.15 | 0.39 | 0.24 | 0.15 | -0.32 | -2.10 |
| **39088** | -0.53 | -0.34 | 0.19 | 0.53 | 0.34 | 0.19 | -0.44 | -2.29 |
| **69421** | -0.53 | -0.38 | 0.15 | 0.53 | 0.38 | 0.15 | -0.46 | -3.03 |
| **97632** | -0.41 | -0.26 | 0.15 | 0.41 | 0.26 | 0.15 | -0.34 | -2.23 |
| **125480** | -0.44 | -0.25 | 0.19 | 0.44 | 0.25 | 0.19 | -0.35 | -1.82 |
| **65185** | -0.34 | -0.12 | 0.22 | 0.34 | 0.12 | 0.22 | -0.23 | -1.05 |
| **38279** | -0.57 | -0.33 | 0.24 | 0.57 | 0.33 | 0.24 | -0.45 | -1.88 |

Compound 123620, 69421, and 97632 appeared to be the strongest electrophiles (among the 11 compounds) with a global hardness of 0.15, however, compound 69421 was the most reactive (Global electrophilicity: −3.03). Of the three representative compounds (139068, 138967, and 38831), compound 139068 and 38831 appeared to be the most reactive (Global electrophilicity: 0.21); however, Compound 138967 was the strongest electrophile (chemical potential: −0.34; HOMO-LUMO gap: 0.18; Global Hardness: 0.18).

Molecular electrostatic potential (MESP) of the lead compounds were calculated; it is the work done in bringing a unit positive charge from infinity to a point [40]; it shows the surface charge distribution on the ligand thereby identify regions that might be involved in the electrophilic reaction (positively charged) and regions involved in the nucleophilic reaction (negatively charged) [40]. The MESP of the representative compounds was visualized and showed positive charge distribution on the ligand (figure 14), this, therefore, confirms that the ligands interact with the BH4 domain via electrophilic reactions.

**Figure 14:**
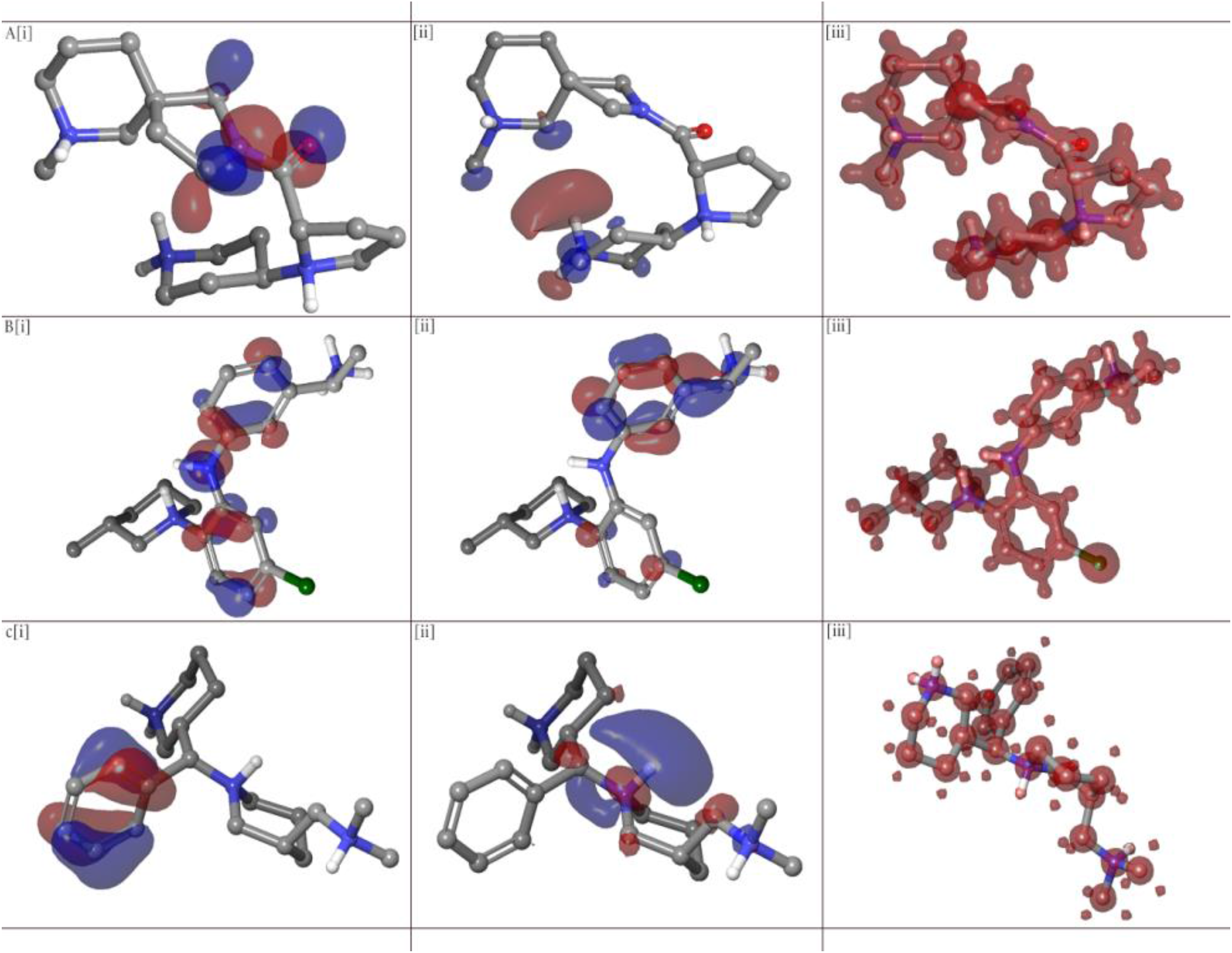
HOMO, LUMO, and MESP for. a) 139068 b) 138967 c) 38831

### 1.6. BCL2 GLU42 mutation analysis

Interaction with non-BH4 Amino acid residue GLU42 has been observed consistently with all the docking protocols and even geometric optimization (QM-MM). We, therefore, sought to investigate if interaction with GLU42 contributes directly to the binding affinity of the compounds. Two mutant BCL2 protein structures were created and minimized: deletion of GLU42 residue (BCL2_Del_) and mutating GLU42 to SER42 (BCL2_SER42_). The result showed binding affinity reduction when docked with the three representative compounds (38831, 138967, 139068) (Table 3). The data suggest that interaction with GLU42 might contribute significantly to the binding of the compounds to the BH4 domain.

**Table 3:** Molecular docking binding affinity with Mutated BCL2 protein.

| Compound ID | BCL2<br>(kcal/mol) | BCL2<sub>Del</sub><br>(kcal/mol) | BCL2<sub>SER42</sub><br>(kcal/mol) |
| --- | --- | --- | --- |
| 38831 | -71.97 | -48.32 | -58.09 |
| 138967 | -62.90 | -57.43 | -44.19 |
| 139068 | -67.22 | -50.28 | -55.93 |

### 1.7. BH4 specificity investigation

One of the downsides of inhibitors that have been developed to targeting the BH3 domain is that the small molecules also bind with other proteins in the BCL2 families (BCL-XL, MCI-1, BFI-1, BCL2A1, etc.) due to the highly conserved nature of the BH3 domains. Since the primary aim of this study is to identify BH4 specific binding small molecules, we, therefore, docked these compounds into the BH3 domain of BCL2, BCL-Xl, and MCL-1 and compare their binding affinity against BDA366.

Using BDA366 as a negative control (since BDA366 has been shown experimentally to be highly selective for BH4), Venetoclax and ABT 737 [8] as a positive control for BCL2 and BCL-XL BH3 docking, and 2,5-substituted benzoic acid [41] as a positive control for MCL-1 BH3 domain docking, the lead compound was docked and binding affinity calculated.

The results showed Compounds 123620 and 69421 had a higher binding affinity (−75.49 and −78.40kcal/mol respectively) for the BCL2 BH3 domain than BDA366 (−70.57kcal/mol) (Table 4) suggesting that this compounds might also bind and interact with BCL2 BH3 domains. However, docking with other BCL2 family proteins none of the compounds bind to the BH3 domain with a binding affinity greater than BDA366 (Table 4).

**Table 4:** BH4 specificity docking.

| Compound ID | BCL2(BH3)<br>(kcal/mol) | BCL-XL(BH3)<br>(kcal/mol) | MCL-1(BH3)<br>(kcal/mol) |
| --- | --- | --- | --- |
| **Venetoclax** | -103.14 | -86.17 | - |
| **ABT737** | -115.44 | -80.24 | - |
| **BDA366** | -70.57 | -55.99 | -84.24 |
| **2,5-substituted benzoic acid** | - | - | -124.33 |
| **38279** | -66.93 | -48.52 | -80.02 |
| **38831** | -70.96 | -43.66 | -62.46 |
| **39088** | -58.2 | -41.09 | -57.68 |
| **65185** | -53.01 | -44.73 | -67.47 |
| **69421** | -78.4 | -47.01 | -58.52 |
| **97632** | -53.25 | -39.65 | -59.28 |
| **123620** | -75.49 | -39.53 | -77.62 |
| **125480** | -62.98 | -40.37 | -69.24 |
| **138967** | -67.68 | -54.66 | -83.84 |
| **139068** | -70.01 | -47.03 | -69.32 |
| **163159** | -67.07 | -44.79 | -55.19 |

## Discussion

BCL2 anti-apoptosis protein has been implicated in several cancer diseases, thus several small molecules have been developed to inhibit its activity. Most of these inhibitors target the BH3 domain of BCL2 however this has not been without its challenges (poor selectivity for BCl2) as none of the small molecule inhibitors is yet to be approved by the FDA [8][10]. Targeting the BH4 domain has shown promising results of converting BCL2 protein form survival to death protein. However, only BDA366 has been reported so far (at the time of writing) as a BH4 specific binding small molecule [30].

Virtually screening ~1,000,000 compounds, 11 compounds showing high binding affinity (−74 to −63kcal/mol) for BCl2 BH4 domain (higher than BDA366: −45kcal/mol) has been identified. In other to elucidate a binding hypothesis for the top 3 compounds, they were subjected to more accurate and rigorous docking simulations: QM-polarized docking (QPLD) and Induced-fit docking (IFD). QPLD investigated the effect of induced charges on the ligands (i.e. on binding interactions and binding affinity) by the active site residues and IFD predicted active site conformational changes that could occur with its effects on ligand binding interactions and binding affinity. A weighted binding affinity average was computed as a way to consolidate the three different binding affinities (i.e. Molecular docking, QPLD, and IFD); the top ranked 3 compounds (139068, 138967, 38831) were chosen as representatives and there corresponding IFD complex were further validated and optimized using QM-MM calculations.

We elucidated a binding hypothesis for the three compounds based on two criteria: firstly consistency of binding interaction throughout all docking protocols and QM-MM validations; secondly new binding interactions with BH4 amino acid residues (AA: 6-31) observed during IFD (we considered IFD to be more accurate since active site conformational changes are put into consideration [33]) and validated in QM-MM calculations. Based on our data we, therefore, suggest the following binding hypothesis: compound 139068: interaction with GLU13, MET16, LYS17, ASP31, and GLU42; compound 138967: interaction with ASP10, ARG12, GLU13, HIS20, MET16, and GLU42; compound 38831: interaction with ASP10, ARG12, GLU13, LYS17, and GLU42.

Works of Han *et al.* [30] suggested the following BH4 residue by which BDA366 regulates BCL2: ASP10, ASN11, ARG12, and GLU13. Also, Hirotani *et al* [9] suggested that interactions with hydrophobic residues of BH4 also regulate the conversion of BCL2 protein from anti-apoptosis to pro-apoptosis protein. Compound 139068 interactions with GLU13 and hydrophobic MET16 suggest that its regulation of BCL2 protein might be via these residues. Compound 138967 and 38831 interaction with ASP10, ARG12, and GLU13 also suggest a plausible mechanism of antagonist activity.

Consistently, interaction with GLU42 was observed among the 3 representative compounds and mostly among the 11 compounds in all docking simulations. We, therefore, investigated the plausible role of GLU42 interactions, as the reason for the generally observed high binding affinity. Mutating BCL2 protein (by deleting GLU42: BCL2_Del_ and mutating to SER42: BCL2_SER42_) and docking the 3 compounds resulted in a fold decrease in binding affinity of the compounds for BCL2 BH4 domain. In confirmation of this observed decrease, recent studies have shown that interaction with GLU42 amino residue plays a significant role in modulating the anti-apoptosis activity of BCL2 protein [42].

Molecular dynamics simulation using normal mode analysis (NMA) showed low eigenvalues for the three compounds which represent high-frequency motions and ease of protein structure deformation (conformational changes). BDA366 binding to BCL2 BH4 domain has been shown to cause a conformational change in BCL2 protein causing its BH3 domain to be “exposed‟ leading to its conversion to pro-apoptosis protein [30]; also, accelerated molecular dynamic (aMD) simulation showed PRO127 and TRP30 rotating to interact (π-π interactions) with BDA366 causing conformation changes in its α helix [43]. These results are consistent with our NMA data which showed BDA366 to have low eigenvalue of 2.682577e-04, this, therefore, suggest that the 3 ligands with similar eigenvalues with BDA366 might cause such conformation changes to BCL2; better still is that the ligands bind with higher binding affinity and with lower eigenvalues (38831 and 139068) than BDA366.

The primary aim of this study is to identify potential small molecules binding to the BCl2 BH4 domain; specificity for this domain was therefore investigated. Using BDA366 as negative control we docked the 11 compounds into the BH3 domains of BCL2 protein itself, MCL-1, and the close relative of BCL-XL. We infer that binding to this BCL2 family protein with a binding affinity lower or equal to BDA366 might indicate specificity for the BH4 domain. Except for compounds 123620 and 69421 having a high binding affinity for BCL2 BH3 domain than BDA366 all other compounds showed equal or lower binding affinity for BCL2 family protein. This might, therefore, suggest a potential preference for the BH4 domain; however, theirs is still a need for experimental validation.

Calculating the electronic properties of the 11 compounds using DFT calculations suggest that the antagonistic activities of these compounds might be via electrophilic reactions. Of the three representative compounds compound, 138967 is suggested as the strongest electrophile however, this did not correspond to it been the highest binding molecule.

## Conclusion

Targeting the BH4 domain in BCL2 protein is a promising strategy in „converting‟ survival proteins to death proteins in cancer cells; identifying and developing BH4 small molecule is therefore important. Using computer simulations, 11 diverse small molecules have been identified; a binding mode for the top ranked 3 compounds have been elucidated and chemical reactivity of the compounds explained. Based on our theoretical data we have suggested these compounds however, experimental data are still needed to validate the antagonistic activity of these compounds.

